# THE IMPACT OF ANKLE IMMOBILITY ON SPRINT CYCLING PERFORMANCE: IMPLICATIONS FOR PARA-CYCLING CLASSIFICATION

**DOI:** 10.64898/2026.05.12.723700

**Authors:** R. Ilse Boot, Ingrid Kouwijzer, Maarten F. Bobbert, Sonja de Groot, Dinant A. Kistemaker

## Abstract

**Purpose:** The para-cycling classification system aims to minimize the impact of impairments on competition outcomes with the help of scientific evidence. This study investigated the impact of unilateral and bilateral ankle immobility on cycling performance, quantified by the maximal average mechanical power output (AMPO) over one revolution relative to that without ankle immobility.

**Methods:** Ten well-trained non-disabled cyclists performed all-out 6-second sprints on a cycle ergometer at 120 rpm under three conditions: without ankle foot orthoses (AFOs), with 1 AFO and with 2 AFOs immobilizing the ankle joint(s). Mechanical power output, pedal forces, cycling kinematics and surface-electromyography were measured. Maximal AMPO; ankle, knee and hip joint AMPO; and the amount of muscle excitation were calculated.

**Results:** With 1 AFO and 2 AFOs, respectively, maximal AMPO was 96% (p<0.05) and 91% (p<0.001) of that without AFOs (1188 W). The decrease in maximal AMPO with ankle immobilization was less than the decrease in ankle joint AMPO (126 W decrease with 2 AFOs; p<0.001), due to an increase in hip joint AMPO (69 W increase with 2 AFOs; p<0.05). The amount of muscle excitation was not significantly different across conditions.

**Conclusions:** These findings provide a first quantitative and mechanistic indication of the impact of ankle immobility on cycling performance, which may offer valuable evidence to support the development of an evidence-based para-cycling classification system.

## Introduction

Para-sports are sports for individuals with physical, vision or intellectual impairments. Over the last decades, para-sports have undergone substantial growth; between the Paralympic Games of Rome in 1960 and Paris in 2024, the number of competing athletes increased from 400 to 4400^1,2^. According to a recent large-scale survey, 80% of the respondents were aware of the Paralympic Games and 41% expressed interest in following them^3^. This makes the Paralympic Games the third most widely recognized sporting event in the world^3^. To enable fair competitions, para-athletes compete in sport-specific classes based on the impact of their impairment on the ability to execute the specific tasks and activities fundamental to the relevant sport^4^. The system that groups para-athletes into classes is called the classification system. For example, the cycling division of para-cycling includes five classes (C1-C5), wherein athletes with musculoskeletal and/or coordination impairments who can safely ride a bicycle compete^5^. Athletes with impairments that are considered to have the most detrimental effect on cycling performance are classified in C1, whereas those with impairments considered to have the least detrimental effect are classified in C5^5^. Each class includes a wide range of eligible impairments (namely: impaired muscle power, impaired passive range of movement, limb deficiency, leg length difference, hypertonia, ataxia and dyskinesia), which may present in different limbs^5^. Hence, large heterogeneity exists in impairments among para-cyclists between and within classes.

The International Paralympic Committee (IPC) mandates the development of evidence-based classification systems for Paralympic sports^4,6^. These classification systems should aim to promote sports participation in athletes with eligible impairments by minimizing the impact of impairments on the outcome of competition. Furthermore, they should be supported by evidence from classification research indicating that this aim has been achieved^4,6^. Specifically, classification research is warranted to investigate the impact of impairments on sports performance. Research for the cycling division of para-cycling is growing^7^, and several methodological approaches have been employed. For example, 1-km track time trial performance of para-cyclists with a unilateral transtibial amputation has been compared to that of para-cyclists with other impairments classified in C4^8^. Another study predicted the impact of a unilateral transtibial amputation on 4-km pursuit performance by modelling both the aerodynamic effects of a prosthesis as well as the reported power asymmetries in cyclists with this impairment^9^. Other work investigated the impact of an impairment on sprint power output by comparing performance between Paralympic track-and-field sprinters with mild to moderate spastic hemiplegic cerebral palsy and non-disabled athletes^10^, or by comparing performance between non-disabled participants with and without the simulated loss of unilateral handgrip^11^. To our knowledge, the biomechanical mechanisms underlying the impact of these and other impairments on cycling performance have only been explored during submaximal cycling (e.g., unilateral transtibial amputation (40-160 W)^12–15^ and single-leg cycling (30-205 W per leg)^16–19^). Despite these efforts, the impact of many eligible impairments on cycling performance and the biomechanical mechanisms underlying this effect remain insufficiently understood. This warrants more research on the impact of impairments on cycling performance.

One impairment that may be particularly relevant for classification in para-cycling is ankle immobility. Recent studies reported an overlap in track time trial performance between C3 and C4 (both men’s 1-km and women’s 500-m^20^), and between C4 and C5 (men’s 1-km^21^). The cause of this overlap is unknown. Although there is large heterogeneity among the impairments within these classes, ankle immobility is among the key determinants for classification in C3 (e.g., complete bilateral ankle immobility), C4 (e.g., complete unilateral ankle immobility), and C5 (e.g., mild to severe unilateral ankle immobility)^5^. The overlap in performance between C3-C4 and C4-C5 may therefore be caused by insufficient knowledge of the impact of ankle immobility on cycling performance. One feasible approach to investigate the impact of ankle immobility on cycling performance would be to compare cycling performance of non-disabled cyclists between wearing and not wearing ankle foot orthoses (AFOs) immobilizing the ankle joint. This approach enables within-participant comparisons of performance, avoiding confounders inherent in comparisons among para-cyclists who represent a small and heterogeneous group in which performance depends not only on impairment but also on factors such as talent, training, and equipment. It provides a controlled setting to investigate the biomechanical consequences of ankle immobilization, which may inform future research employing approaches with greater external validity (e.g., studies including para-cyclists with ankle immobility in a competitive setting). A valid determinant for (para-)cycling performance is the maximal average mechanical power output (AMPO) over one full revolution^22^. The maximal AMPO with ankle immobility expressed as a percentage of that without ankle immobility is then an objective measure for the impact of ankle immobility on (maximal) cycling performance. To our knowledge, previous experimental studies involving non-disabled participants have only investigated the impact of wearing AFOs on cycling performance during submaximal recumbent cycling (17-94 W)^23–25^. All in all, this warrants investigating the impact of ankle immobility on cycling performance in the context of para-cycling.

Here, the primary aim was to investigate the impact of unilateral and bilateral ankle immobility on cycling performance, quantified by the maximal AMPO relative to that without ankle immobility. For this, well-trained non-disabled cyclists performed all-out sprints on a cycle ergometer with and without AFOs immobilizing the ankle joint. The secondary aim was to explore potential underlying mechanisms of the observed impact of ankle immobility on maximal AMPO. This included an inverse dynamics analysis to determine the impact of ankle immobility on the AMPO around the ankle, knee and hip joints. In addition, surface-electromyography (EMG) was used to determine the impact of ankle immobility on the amount of muscle excitation.

## Materials and methods

### Participants

Ten well-trained non-disabled cyclists participated in this study. Inclusion criteria were: aged 18-65 years; self-reported 1-second sprint power output ≥1000 W; European shoe size ≥42. Candidates were excluded if they had any musculoskeletal, cardiovascular and/or pulmonary disease or injury affecting cycling performance. Participants were medically screened using the Physical Activity Readiness Questionnaire (PAR-Q)^26^. All participants were male and had a median (Q1-Q3) age of 28 (24-38) years, body mass of 79 (71-94) kg, body height of 1.82 (1.80-1.86) m, and shoe size of 44 (42-45). Participants had 10 (4-15) years of cycling experience and trained 6 (3-12) hours for cycling weekly. The right leg was the dominant leg for all but one participant. Cycling levels ranged from recreational (N=5) to regional competition (N=3), national competition (N=1), and international competition (N=1). The participants were instructed to avoid strenuous activity for 24 hours and alcohol for 4 hours preceding the experiment. The experimental procedures were reviewed and approved by the Scientific and Ethical Review Board of the Faculty of Behavioural & Movement Sciences, Vrije Universiteit Amsterdam. Participants were informed about the experimental procedures and gave written informed consent before taking part in the experiments.

### Procedures

The participants performed all-out sprints on a racing bicycle mounted on a Cyclus2 ergometer (RBM elektronik-automation GmbH, Leipzig, Germany; see Fig. 1A) under three conditions: 1) without AFOs, 2) with 1 AFO on the dominant ankle, and 3) with 2 AFOs, one on each ankle. We used custom-made AFOs that were specifically designed for high stiffness while remaining lightweight (De Hoogstraat Orthopedietechniek, Utrecht, the Netherlands). These AFOs fixed the ankle joint in neutral position (sole of foot in 90° relative to shank). We chose a neutral position, because 1) it is likely the most common representation by para-cyclists, and 2) it avoids confounders related to unnecessary alterations in cycling kinematics, as it is close to the previously reported average ankle joint angle when cycling with ankle mobility^27^. If possible, participants wore the AFO in their own cycling shoe. Otherwise, participants wore a cycling shoe provided by the experimenter with either size 45 or 46.5. When not wearing an AFO, participants wore their own shoe. For both shoes, adjusting the cleat position to the participant’s preference was allowed. Ergometer settings were kept constant across conditions and were set to values (close to) optimal in sprint cycling. Specifically, cadence was set to 120 rpm^28,29^ and saddle height was set such that the minimal knee joint angle (averaged over ten revolutions) equalled 35° in the 2 AFOs condition^30,31^ (see Fig. 1B for knee joint angle definition). Seat tube angle and handlebar position were kept to the standard values for the bicycle (see Fig. 1A).

**Figure 1.**
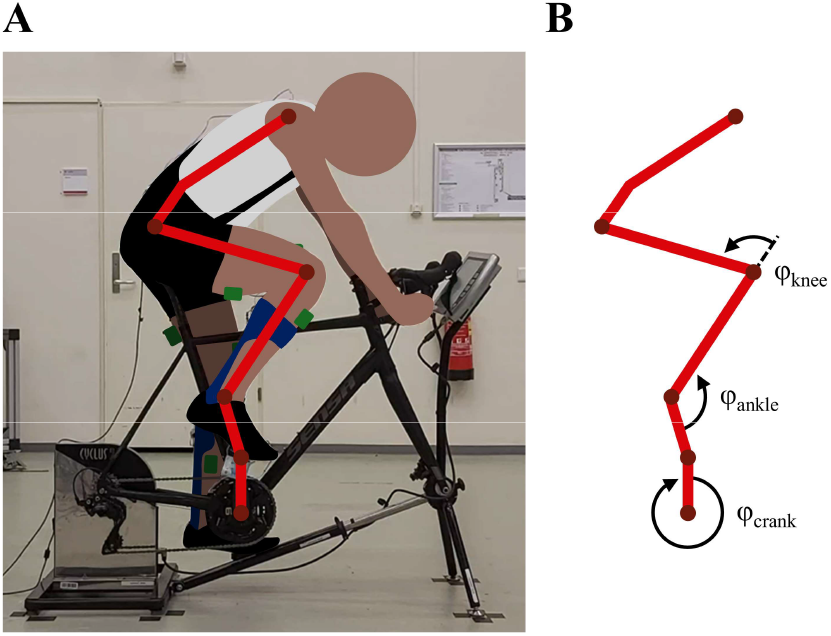
(A) Experimental setup. The participant cycles on a bicycle mounted on a Cyclus2 ergometer while wearing ankle foot orthoses (AFOs; indicated in blue). Mechanical power output and pedal forces were measured by the pedals. Cycling kinematics were determined using an active motion capture system (indicated by red circles; see text for detailed marker placement). Muscle excitations were determined using a wireless EMG system (indicated by green rectangles; see text for detailed electrode placement). (B) Segment model with joint angle definitions (left side not shown). Top dead centre (TDC) is defined as a crank angle of 0°, which is the position indicated in the figure. From bottom bracket to shoulder, the segment model comprised a crank, foot, shank, thigh, and trunk (incl. kink for visualization). EMG, electromyography; φ_crank_, crank angle; φ_ankle_, ankle joint angle; φ_knee_, knee joint angle.

The experiment comprised two sessions, separated by one week (see Fig. 2). During the first session, the participants familiarized with the experimental setup during which they sprinted in the three conditions. During the second session, the measurements were performed. Each session consisted of three blocks of sprints, one condition per block, separated by 5 minutes of rest. The order of conditions was counter-balanced (across participants) in the first session. In the second session, this order was reversed. Each block started with a warm-up in the corresponding condition, which also functioned as familiarization. This warm-up consisted of 7 minutes of cycling at 100 W, which was directly followed by 3 minutes of cycling at 150 W. After 2 minutes of rest, the participants performed 6-second isokinetic sprints. Sprints were performed using the Cyclus2’s preprogrammed ‘isokinetic maximum strength test’. During this test, the ergometer modified the pedal resistance such that the cadence equalled 120 rpm irrespective of the power output by the participant. For proper functioning, the test required selecting an ‘initial load’ of the ergometer. During the first session, this initial load was tuned for each participant and condition with an accuracy of 50 N. For familiarization and proper tuning of the initial load, three sprints per condition were performed in the first session. To minimize fatigue, we limited the number of sprints to two in the second session. The participants were instructed to slowly increase their cadence towards 90 rpm (which took about ten revolutions). When 90 rpm was reached, the 6-second isokinetic sprint started. Participants were given verbal encouragement and were instructed to remain seated.

**Figure 2.**
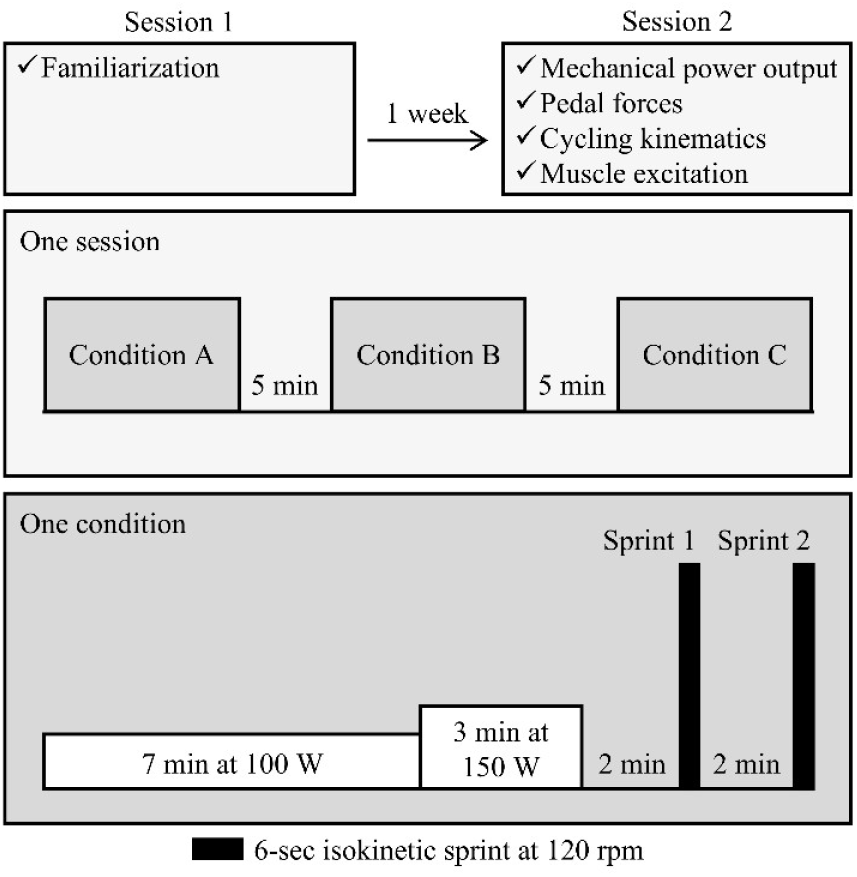
Experimental protocol. The protocol comprised two sessions, each consisting of sprints in all conditions: 1) without AFOs, 2) with 1 AFO on the dominant ankle, and 3) with 2 AFOs. The order of conditions was counter-balanced across participants, resulting in an A-B-C sequence. In the first session, three sprints per condition were performed (third sprint not shown). In the second session, two sprints per condition were performed. Sprints were 6-second isokinetic sprints at 120 rpm. Vertical axes denote power output. Horizontal axes denote cycling duration. AFO, ankle foot orthosis.

### Data acquisition

We used SRM pedals to measure crank angle, crank angular velocity, and tangential and radial pedal forces at 200 Hz (SRM X-Power Road; Schoberer Rad Meßtechnik, Jülich, Germany). We validated these pedals and found that the pedal forces demonstrated good static validity.

Furthermore, the mechanical power output determined from these pedals showed good dynamic validity, which was comparable to or better than that of two other power meters (see Supplemental Digital Content, Validation power meters).

To determine the three-dimensional kinematics (at 100 Hz) of the shoulder joint, hip joint, knee joint, ankle joint, pedal axis, and bottom bracket, an Optotrak motion capture system was used (Northern Digital Inc., Ontario, Canada). For this, active markers were mounted on the lateral side of acromion, trochanter major, epicondylus lateralis femoris, and malleolus lateralis. Since the shoe occluded the pedal axis, we reconstructed the position of the pedal axis from the measured position of two markers mounted on a lightweight foam block that was attached to the bottom of the pedal. The position of the bottom bracket was then reconstructed from the position of the pedal axis.

Bipolar surface-electromyography (EMG) from four muscles (m. vastus medialis (VAS), m. biceps femoris caput longum (HAMb), m. tibialis anterior (TIB), and m. soleus (SOL)) was measured at 2148 Hz using a Delsys Trigno wireless EMG system (Delsys Inc., Natick, MA, USA). Localization of the muscles of interest, preparation of the skin and placement of the EMG electrodes were conducted following SENIAM guidelines^32^. The Delsys Trigger Module ensured time-synchronization between the motion capture and EMG systems. All measurements were performed bilaterally (see Fig. 1A).

### Data analysis

Data analysis was performed using custom-written programs in MATLAB (version: R2024a; MathWorks, Natick, MA, USA). From the measured three-dimensional kinematics, those in the sagittal plane were calculated and used for further analyses. In this plane, the crank angle and crank angular velocity were calculated. The crank angle was used to time-synchronize the data from the pedals with those of the motion capture system. Since the Optotrak motion capture system is very accurate (precision: 0.125 mm)^33^, the mechanical power output was calculated using the crank angular velocity from the motion capture system. Similarly, the pedal force vectors were calculated using the crank angle from the motion capture system. Individual revolutions were identified that started and ended when the crank of the dominant leg was at top dead centre (TDC, here defined as a crank angle of 0°; position indicated in Fig. 1B). For each sprint, the three revolutions with the highest AMPO were selected. Overall, this resulted in data of six revolutions for each participant and condition.

A segment model was created for each revolution (see Fig. 1B). For this, the segment lengths and angles were fitted to the experimentally observed kinematic data. During an all-out sprint, motion of the hip joint position relative to the saddle is difficult to completely avoid. To prevent this (spurious) motion from affecting the estimated position of particularly the ankle joint, we created the segment model using a stepwise optimization. Here, we first optimized the segment length and angles of the foot (pedal axis-ankle) such that the root-mean-squared difference between the calculated and measured position of the ankle joint was minimized. Then, we optimized the segment length and angles of the shank (ankle-knee), given the calculated position of the ankle joint. Under the assumption that the participants indeed were capable of remaining seated, we then optimized the segment length and angles of the thigh (knee-hip), such that the root-mean-squared difference between the calculated position of the hip joint and the measured average position of the hip joint (one value) was minimized. Finally, we optimized the segment length and angles of the trunk (hip-shoulder), given the calculated average position of the left and right hip joints (value changing over time). An inverse dynamics analysis was performed using the segment model and pedal force vectors. The relative segment parameters were taken from previous research and are shown in Table 1^29,34,35^.

**Table 1.** Relative segment parameters.

|  | **Crank** | **Foot** | **Shank** | **Thigh** |
| --- | --- | --- | --- | --- |
| **Segment mass** | 0.2000<sup>†</sup> | 0.0146 | 0.0418 | 0.1000 |
| relative to body mass [kg] |  |  |  |  |
| **Location of COM (distal to proximal)** | 0.5000 | 0.7273 | 0.5677 | 0.5669 |
| relative to segment length [m] |  |  |  |  |
| **Radius of gyration relative to COM** | 0.4159 | 0.5456 | 0.3026 | 0.3238 |
| relative to segment length [m] and segment mass [kg] |  |  |  |  |

Relative segment parameters used for the inverse dynamics analysis. Values were adopted from previous research<sup>29,34,35</sup>. COM, centre of mass; <sup>†</sup>crank mass is not relative to body mass.

The amount of muscle excitation was determined from the EMG signals^36^. For this, the EMG signals were band-pass filtered (zero-phase IIR filter, order: 2, cut-off frequencies: 20 and 500 Hz), rectified and low-pass filtered (zero-phase IIR filter, order: 2, cut-off frequency: 15 Hz) to obtain EMG envelopes. The EMG envelopes were normalized in several steps. First, for each of the eight muscles per participant, the EMG envelopes for the condition without AFOs were averaged across the six selected revolutions. Second, the averaged EMG envelopes were smoothed using a three-sample moving mean. Third, the original EMG envelopes for each revolution and condition were normalized to the maximum value of the smoothed EMG envelope for the corresponding muscle and participant. To minimize the effect of noise, values in the normalized EMG envelopes that fell below 10% were set to zero. The amount of muscle excitation was quantified by the integral of the resulting normalized EMG envelopes with respect to the crank angle.

Within participants, all outcome measures were averaged across the six selected revolutions per condition. For example, the reported maximal AMPO for individual participants equals the average across these six revolutions. The relative maximal AMPO was calculated by expressing the maximal AMPO of the conditions with 1 AFO and 2 AFOs as a percentage of that without AFOs. To investigate potential underlying mechanisms of the observed relative maximal AMPO, we also analysed the relative AMPO around the ankle, knee and hip joints (derived from the inverse dynamics analysis) as well as the relative amount of muscle excitation. Across participants, we report medians and interquartile ranges (Q1-Q3) in accordance with the performed non-parametric statistics (see Statistics). Time-varying variables are exceptions, as for these variables the mean and standard error of the mean (SEM) across participants are plotted against crank angle.

### Statistics

Statistical tests were performed in MATLAB. Friedman’s analysis of variance (ANOVA) was used to test whether maximal AMPO, joint AMPO, and amount of muscle excitation significantly differed (p<0.05) across the three conditions. If significant, post-hoc pairwise comparisons with Tukey-Kramer correction were performed. To check whether the imposed cadence was indeed similar across conditions, we additionally tested for statistically significant differences in cadence (as a function of crank angle) among the three conditions using a statistical non-parametric mapping (SnPM) ANOVA^37^.

## Results

Before exploring the impact of wearing AFOs on maximal AMPO, we checked whether our experimental protocol was performed successfully. The AFOs greatly reduced ankle mobility, as the ankle joint range of motion (ROM) with AFO was 33 (24-45)% of that without AFO (with AFO: 13 (12-15)°; without AFO: 38 (33-44)°). Immobilizing the ankle joint in neutral position successfully resulted in a similar average ankle joint angle with as without AFO: the immobilized ankle joint was 4° more in dorsiflexion (5° more in dorsiflexion – 2° more in plantarflexion). Horizontal saddle position (which corresponds to the hip joint position in the segment model) was 0.21 (0.20-0.23) m posterior from the bottom bracket. Vertical saddle position was 0.81 (0.79-0.84) m above the bottom bracket. The minimal knee joint angle with AFO was 38 (33-44)°, which is close to the intended 35°. Cadence (as a function of crank angle) did not differ significantly across conditions (see Fig. 3), and averaged 119 (119-120) rpm. Fig. 4 displays the tangential and radial pedal forces of the dominant leg for each condition. Animations of the cycling kinematics, pedal forces and muscle excitations can be found online (see Supplemental Digital Content; Animation of cycling kinematics, pedal forces and muscle excitations).

**Figure 3.**
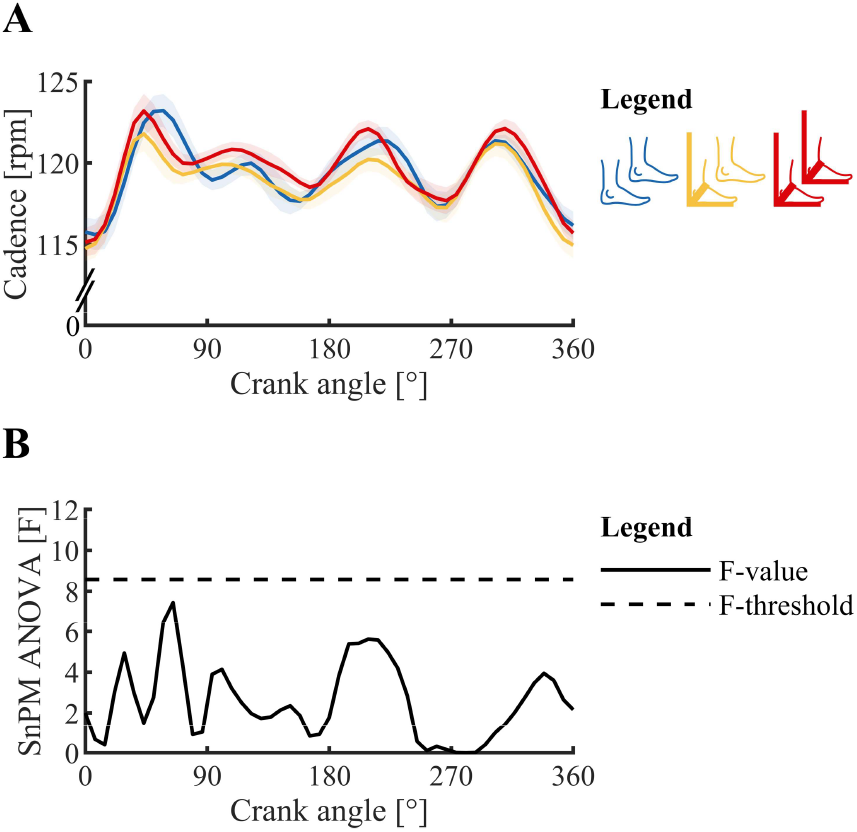
(A) Cadence against crank angle for each condition. Lines with contours represent the mean and standard error of the mean (SEM) across participants. (B) The F-value from the statistical non-parametric mapping (SnPM) ANOVA against crank angle. As the F-value did not reach the threshold throughout the revolution, cadence did not differ significantly across conditions.

**Figure 4.**
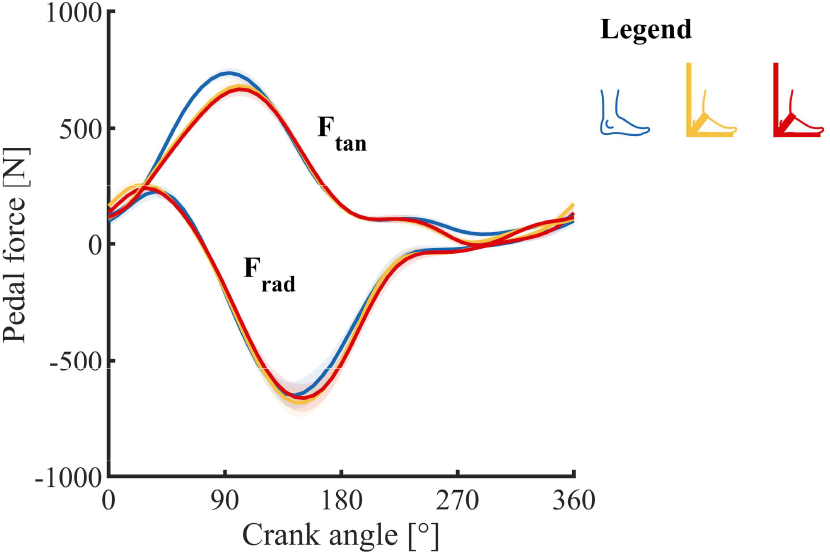
Tangential (F_tan_) and radial (F_rad_) pedal forces of the dominant leg against crank angle. Lines with contours represent the mean and standard error of the mean (SEM) across participants.

### Maximal AMPO

The mechanical power output is shown in Fig. 5A (Total). Fig. 5B displays the corresponding relative maximal AMPO (summed for the dominant and non-dominant legs). Maximal AMPO significantly differed across conditions, and was significantly lower with both 1 AFO and 2 AFOs than without AFOs (see Table 2). Relative maximal AMPO with 1 AFO and 2 AFOs was 96 (92-97)% and 91 (89-94)%, respectively.

**Table 2.** Statistical findings.

|  | Median (Q1-Q3) |  |  | Friedman's ANOVA [p-values] | Post-hoc pairwise comparisons [p-values] |  |  |
| --- | --- | --- | --- | --- | --- | --- | --- |
|  | 0 AFOs | 1 AFO | 2 AFOs |  | 0 AFOs - 1 AFO | 0 AFOs - 2 AFOs | 1 AFO - 2 AFOs |
| **AMPO [W]** |  |  |  |  |  |  |  |
| Maximal AMPO | 1188<br>(1028-1260) | 1151<br>(956-1200) | 1083<br>(959-1165) | **<0.001** | **0.04** | **<0.001** | 0.17 |
| Ankle joint AMPO | 163<br>(142-206) | 97<br>(77-118) | 37<br>(22-48) | **<0.001** | 0.07 | **<0.001** | 0.07 |
| Knee joint AMPO | 486<br>(442-515) | 470<br>(383-553) | 455<br>(385-523) | 0.67 |  |  |  |
| Hip joint AMPO | 464<br>(395-562) | 477<br>(421-586) | 533<br>(460-631) | **0.01** | 0.50 | **0.01** | 0.17 |
| **Amount of muscle excitation [%]** |  |  |  |  |  |  |  |
| VAS | 100 | 103<br>(94-105) | 106<br>(102-107) | 0.12 |  |  |  |
| HAMb | 100 | 96<br>(89-107) | 97<br>(88-107) | 0.90 |  |  |  |
| TIB | 100 | 93<br>(74-101) | 87<br>(82-108) | 0.50 |  |  |  |
| SOL | 100 | 97<br>(83-115) | 84<br>(69-88) | 0.08 |  |  |  |

Outcomes of the statistical tests performed. Bold p-values denote statistical significance ( $p < 0.05$ ). AFO, ankle foot orthosis; AMPO, average mechanical power output; VAS, m. vastus medialis; HAMb, m. biceps femoris caput longum; TIB, m. tibialis anterior; SOL, m. soleus.

**Figure 5.**
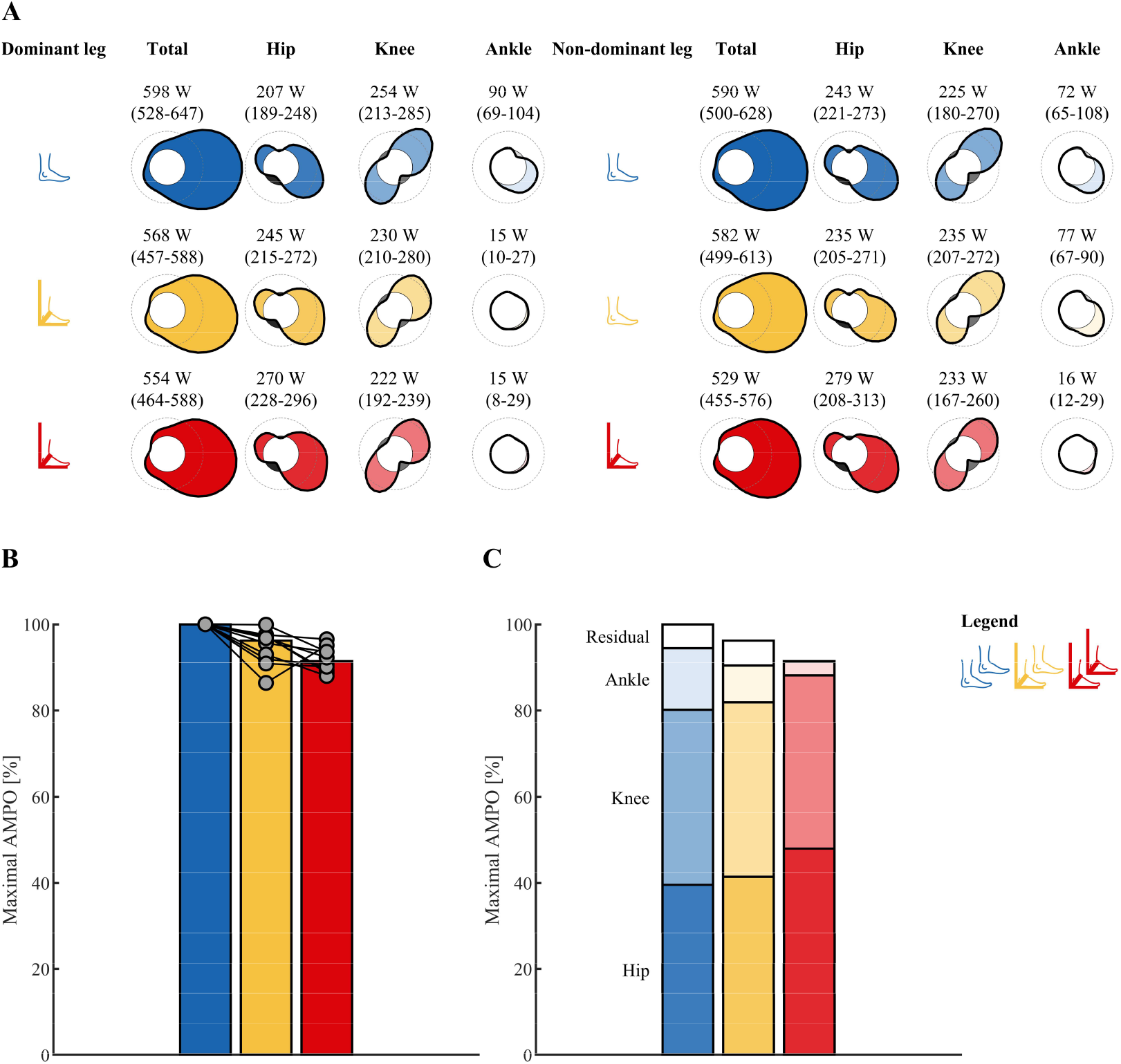
The impact of ankle immobility on maximal AMPO and joint AMPO. (A) The mechanical power output measured by the pedals (Total) and calculated around the ankle, knee and hip joints against crank angle for both the dominant and non-dominant legs during each condition. Thin black circles represent 0 W. Dashed grey circles represent 500 W. Bold black lines represent the mean mechanical power output across participants; coloured fillings represent positive power, greyed fillings represent negative power. Values indicate the median (Q1-Q3) AMPO across participants. (B) Maximal AMPO relative to that without AFOs (summed for the dominant and non-dominant legs). Bars represent median values across participants. Individual participants are indicated by the circles connected with lines. (C) The relative contributions of the ankle, knee and hip joints (summed for the dominant and non-dominant legs) to maximal AMPO. Bars represent median values across participants. Residual indicates the AMPO not explained by the AMPO around the three joints. AMPO, average mechanical power output.

### AMPO around ankle, knee and hip joints

The mechanical power output around the ankle, knee and hip joints is shown in Fig. 5A. Fig. 5C displays the relative contributions of the ankle, knee and hip joints (summed for the dominant and non-dominant legs) to maximal AMPO. Maximal AMPO is not equal to the sum of the joint AMPOs; this discrepancy is indicated by the residual term. This residual is the result of assumptions in our inverse dynamics analysis (e.g., rigid segments, the hip being fixed to the saddle, and no power generated by the upper body) as well as the use of median rather than mean joint AMPO values. (Note that, by definition, instantaneous total power does not equal the sum of the instantaneous joint powers, due to changes in kinetic and potential energy^34^. This discrepancy, however, cancels out when averaged over a full revolution; in periodic movements, the net change in kinetic and potential energy over one cycle is zero.) The AMPO around the ankle joint significantly differed across conditions, and was found to be significantly lower with 2 AFOs than without AFOs (see Table 2). Ankle joint AMPO with 2 AFOs was only 19 (16-28)% of that without AFOs, which is a direct consequence of ankle immobilization. The AMPO around the knee joint did not significantly differ across conditions (see Table 2). The AMPO around the hip joint did significantly differ across conditions, and was found to be significantly higher with 2 AFOs than without AFOs (see Table 2). Hip joint AMPO with 2 AFOs was 112 (107-127)% of that without AFOs. Due to this increase in AMPO around the hip joint, the decrease in maximal AMPO with wearing AFOs was less than the decrease in AMPO around the ankle joint.

### Amount of muscle excitation

Fig. 6 shows the normalized EMG envelopes (A) and the corresponding amount of muscle excitation (summed for the dominant and non-dominant legs) relative to that without AFOs (B). No significant differences in the amount of muscle excitation across conditions were found (see Table 2). This was also true when we used normalized EMG envelopes without cut-off at 10% or non-normalized EMG envelopes without cut-off (results not shown).

**Figure 6.**
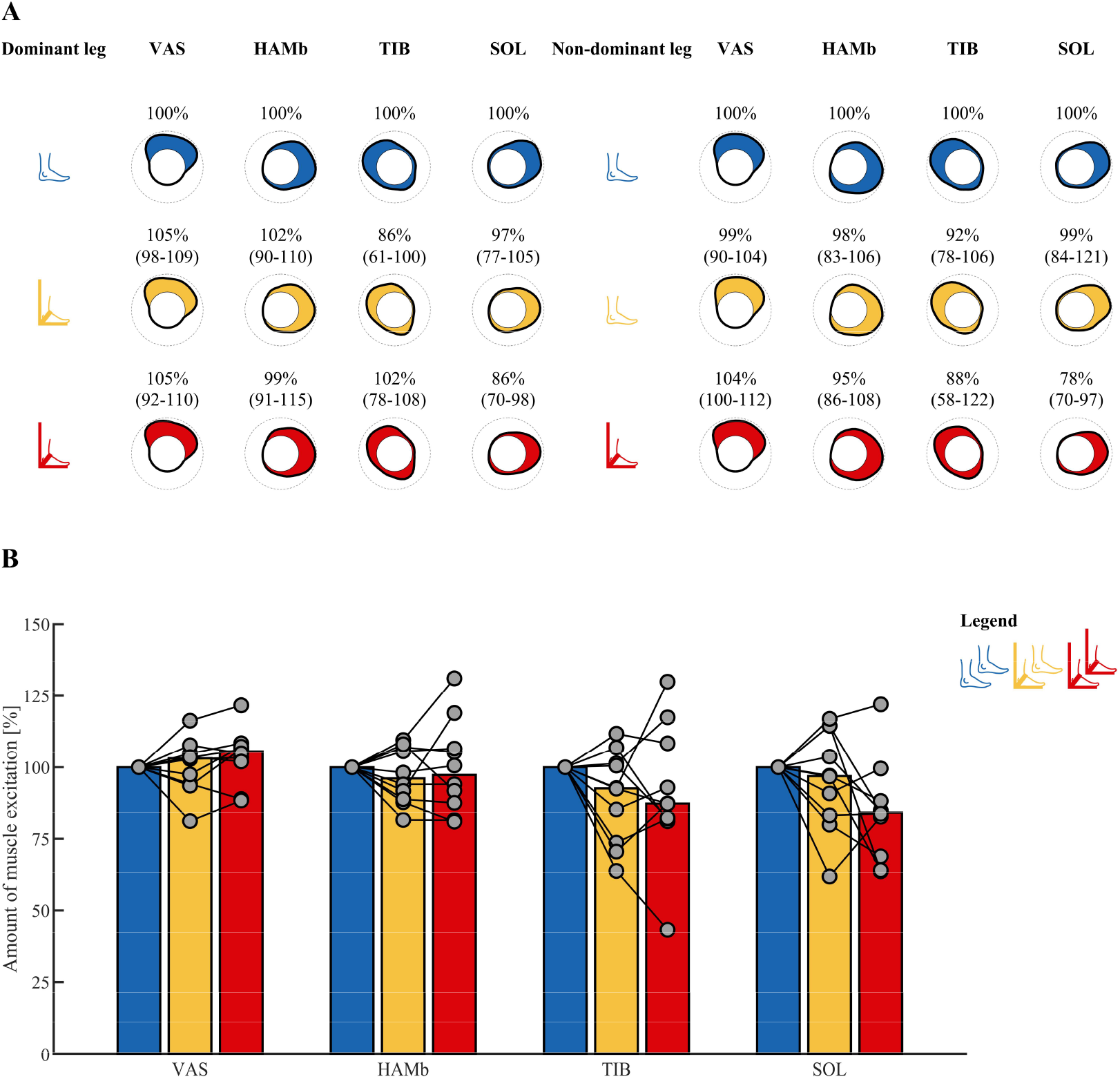
The impact of ankle immobility on the amount of muscle excitation. (A) The muscle excitation (represented as the normalized EMG envelopes) for VAS, HAMb, TIB and SOL for both the dominant and non-dominant legs during each condition. Thin black circles represent 0% muscle excitation. Dashed grey circles represent 100% muscle excitation. Bold black lines with coloured fillings represent the mean muscle excitation across participants. Values indicate the median (Q1-Q3) muscle excitation across participants. (B) The amount of muscle excitation (summed for the dominant and non-dominant legs). Bars represent median values across participants. Individual participants are indicated by the circles connected with lines. EMG, electromyography; VAS, m. vastus medialis; HAMb, m. biceps femoris caput longum; TIB, m. tibialis anterior; SOL, m. soleus.

## Discussion

The primary aim of this study was to investigate the impact of unilateral and bilateral ankle immobility on cycling performance, quantified by the maximal AMPO relative to that without ankle immobility. We found that the relative maximal AMPO with 1 AFO and 2 AFOs was 96% and 91% of that without AFOs, respectively. The secondary aim was to explore potential underlying mechanisms of the observed impact of ankle immobility on maximal AMPO. We found that the decrease in maximal AMPO with wearing AFOs was less than the decrease in AMPO around the ankle joint, due to an increase in AMPO around the hip joint. The amount of muscle excitation did not differ significantly across conditions. In the next sections, we will first discuss whether it was feasible to use AFOs to simulate an ankle mobility impairment. Once this is confirmed, we will explore the mechanisms that may underlie the observed impact of ankle immobility on cycling performance. We finish this discussion with the implications of our findings for the para-cycling classification system.

### Simulation of ankle immobility using AFOs

Using AFOs to immobilize the ankle joint of non-disabled cyclists enabled within-participant comparisons. This avoided confounders inherent in between-participant comparisons and enabled investigating the impact of ankle immobility objectively by expressing the maximal AMPO with ankle immobility as a percentage of that without ankle immobility. Crucial for this approach is that the AFOs were capable of adequately simulating an ankle mobility impairment. We found that the AFOs greatly reduced ankle mobility; the ankle joint ROM with AFO was reduced to 13°, compared to that without AFO of 38°. Hence, the AFOs successfully simulated an ankle mobility impairment.

Immobilizing the ankle joint will unavoidably alter the cycling kinematics, and thereby the muscle-tendon-complex (MTC) length and velocity^38^. As the power generated by muscles depends on the MTC length and velocity^39^, these alterations in cycling kinematics alone will impact cycling performance. To prevent the impact of wearing AFOs on cycling performance from being dominated by additional unnecessary alterations in cycling kinematics, we aimed to minimize the differences in cycling kinematics beyond those unavoidable with ankle immobilization by 1) immobilizing the ankle joint in the average ankle joint angle with ankle mobility and 2) imposing isokinetic cycling at 120 rpm. Previous literature has reported that during sprint cycling at 110 rpm the ankle joint is on average in neutral position^27^. Indeed, we found that immobilizing the ankle joint in neutral position resulted in the ankle joint angle with AFO being similar to the average ankle joint angle without AFO. Cadence (as a function of crank angle) did not differ significantly across conditions, showing similar magnitudes and trends throughout the revolution. We therefore argue that the impact of wearing AFOs on cycling performance was not dominated by unnecessary alterations in cycling kinematics. Hence, we are confident that the decrease in maximal AMPO observed when wearing AFOs was specifically due to ankle immobility.

### Mechanisms underlying the impact of ankle immobility on cycling performance

Having confirmed that the AFOs successfully simulated an ankle mobility impairment, our findings indicate that the decrease in maximal AMPO with ankle immobilization was less than the decrease in AMPO around the ankle joint, due to an increase in AMPO around the hip joint. Explaining the drastic decrease in AMPO around the ankle joint is straightforward: as the ankle joint angular velocity with AFO is almost zero, the AMPO around the ankle joint is almost zero as well. On the level of the muscles, the MTC velocity of the mono-articular ankle joint muscles is almost zero, resulting in an almost zero AMPO generated by these muscles. We found trends of a lower amount of excitation for TIB and SOL. Presumably, participants decreased the excitation of these muscles, because these muscles contributed little to the maximal AMPO with ankle immobilization. By contrast, explaining the increase in AMPO around the hip joint is complicated. This increase should be the result of an increase in AMPO of one or more muscles. Note that these muscles are not necessarily those crossing the hip joint, as bi-articular muscles affect the distribution of muscle power among the joints in an inverse dynamics analysis^22^. An increase in muscle AMPO can be the result of 1) changes in MTC length and velocity, and changes in muscle excitation across conditions. For the first, cycling kinematics differed slightly across conditions, inevitably leading to small differences in MTC length and velocity. Unfortunately, as these differences in MTC length and velocity cannot be assessed experimentally, it is not possible to estimate differences in muscle AMPO due to changes in cycling kinematics. Our EMG analysis does not provide evidence for the second. We showed that the amount of muscle excitation for VAS and HAMb was not significantly different across conditions. This is in contrast to the increased muscle excitation hypothesized by a forward dynamics musculoskeletal modelling study investigating the impact of ankle immobility on maximal AMPO during functional electrical stimulation (FES) cycling^40^. Perhaps, our EMG data were not sensitive enough to detect the differences in excitation leading to a 12% increase in hip joint AMPO. Alternatively, changes in muscle excitation may have occurred in muscles other than the ones we were able to record EMG data from. Overall, it remains unclear which muscle(s) increased their AMPO upon ankle immobilization and by what mechanism. A forward dynamics musculoskeletal modelling approach in the context of para-cycling may be warranted to explain the observed increase in hip joint AMPO.

### Ankle immobility in para-cycling classification

Along with multiple heterogeneous impairments, complete unilateral and bilateral ankle immobility are currently classified in C4 and C3, respectively^5^. In this context, our findings may be slight overestimates; the AFOs simulated incomplete ankle immobility in non-disabled cyclists. We may partly account for this potential overestimate by subtracting the AMPO delivered around the immobilized ankle joint(s) from the maximal AMPO, which reduced the relative maximal AMPO from 96% to 94% for 1 AFO and from 91% to 88% for 2 AFOs. Still, these percentages do not fully represent a complete ankle mobility impairment: for ankle immobility to be classified as complete, calf muscle weakness must be present (personal communication)^5^. The participants in the present study did not exhibit calf muscle weakness and could presumably generate more power with m. gastrocnemius (GAS) than para-cyclists. Unfortunately, we cannot account for this difference, as the experimental data do not enable us to determine the AMPO delivered by individual muscles. To address this, a forward dynamics musculoskeletal modelling approach may be warranted to simulate the impact of complete ankle immobility with and without calf muscle weakness on maximal AMPO. Nevertheless, it seems unlikely that accounting for the AMPO generated by GAS alone would largely decrease the relative maximal AMPO. Thus, relative maximal AMPO with complete unilateral and bilateral ankle immobility may be close to 94% and 88%, respectively. These percentages, together with the observed increase in hip joint AMPO, provide a first quantitative and mechanistic indication of the impact of ankle immobility on cycling performance and may inform the development of an evidence-based para-cycling classification system.

### Perspective

At present, it remains unknown how the observed relative maximal AMPO for unilateral and bilateral ankle immobility compares to that of other impairments eligible for para-cycling. Using a similar experimental approach, future research may therefore investigate the impact of additional impairments on maximal AMPO. In addition, future research may verify our findings in para-cyclists with ankle immobility, as the impact of an impairment on cycling performance of para-cyclists in a competitive setting could differ from the impact of a simulated impairment in well-trained non-disabled cyclists in an experimental setting. Furthermore, future research may employ a forward dynamics musculoskeletal modelling approach to provide further insight into the mechanisms underlying the observed impact of ankle immobility on maximal AMPO. Based on the current findings in well-trained non-disabled cyclists, sprint cycling performance with unilateral and bilateral ankle immobility reaches 96% and 91% of that without ankle immobility, respectively.

## Supporting information

Validation power meters

Animation of cycling kinematics, pedal forces and muscle excitations

## Acknowledgements

This research was conducted with support from the Union Cycliste Internationale (UCI). The funding body was not involved in the study’s design, data analysis, interpretation of the findings, or in the writing and publication of this manuscript.

## Data availability statement

All raw and processed data, together with the analysis code, are available in the Zenodo repository: doi:10.5281/zenodo.20080618

