## Supplementary material for "THE IMPACT OF ANKLE IMMOBILITY ON SPRINT CYCLING PERFORMANCE: IMPLICATIONS FOR PARA-CYCLING CLASSIFICATION": Validation power meters

In the main article, mechanical power output was measured using SRM pedals (SRM X-Power Road; Schoberer Rad Meßtechnik, Jülich, Germany). In this supplementary, we determined the static validity of the pedal forces and the dynamic validity of the mechanical power output measured by these pedals. Where possible, the dynamic validity was compared to the mechanical power output measured by an SRM crank (SRM Science Track; Schoberer Rad Meßtechnik, Jülich, Germany) and a Cyclus2 ergometer (RBM elektronik-automation GmbH, Leipzig, Germany).

*Static validation*

We validated the tangential and radial pedal forces measured by the pedals statically. For this, we placed a bicycle either upright or upside down on a platform and hung weights on the left or right pedal at different crank angles (see Table S1). One experimenter held the bicycle and prevented the crank from rotating by a combination of: placing the front wheel against a wall (when the bicycle was upright); holding the rear wheel (when the bicycle was upside down); pushing the brakes; and applying force to the other pedal.

Retrieving pedal force data at 200 Hz is only possible when the crank is rotating. Therefore, we were limited to directly reading the tangential and radial pedal force magnitudes from the X-Power iOS app. To determine the crank angle, we mounted five markers of an Optotrak motion capture system on the bottom bracket, pedal axis and on three locations on the frame of the bicycle (Northern Digital Inc., Ontario, Canada). We converted the measured three-dimensional marker positions to the corresponding marker positions in a

**Table S1.** Experimental setup for statically validating the pedal forces measured by the pedals

| **Pedal** | **Left** | | | | **Right** | | | |
|---|---|---|---|---|---|---|---|---|
| **Bicycle orientation** | **Upright** | | **Upside down** | | **Upright** | | **Upside down** | |
| **Mass [kg]** | **19.612** | **39.674** | **19.612** | **39.674** | **19.612** | **39.674** | **19.612** | **39.674** |
| **Crank angle [°]** | 9 | 43 | 200 | - | 55 | 55 | 234 | - |
| | 57 | 79 | 247 | | 126 | 113 | 295 | |
| | 123 | 128 | 269 | | 174 | 177 | 343 | |
| | 158 | | | | | | | |
| | 320 | | | | | | | |

The different combinations of pedal, bicycle orientation, mass, and crank angle used for statically validating the pedal forces measured by the pedals. Crank angle is defined as in Fig. 1B.

two-dimensional coordinate system of the bicycle: one axis in the driving direction and one in the vertical direction. Once we determined the crank angle in this two-dimensional coordinate system, we calculated the imposed tangential and radial pedal force magnitudes. We validated the measured with the imposed tangential and radial pedal force magnitudes by calculating the bias, precision and root-mean-squared error (RMSE).

Fig. S1 shows the measured and imposed tangential and radial pedal force vectors for each crank angle. In Fig. S2, the measured and imposed tangential and radial pedal force magnitudes are compared. The bias ± precision (RMSE) for the tangential and radial pedal force magnitudes were -1 ± 5 N (5 N) and -4 ± 6 N (7 N), respectively. These results indicate that the pedal forces demonstrate good static validity.

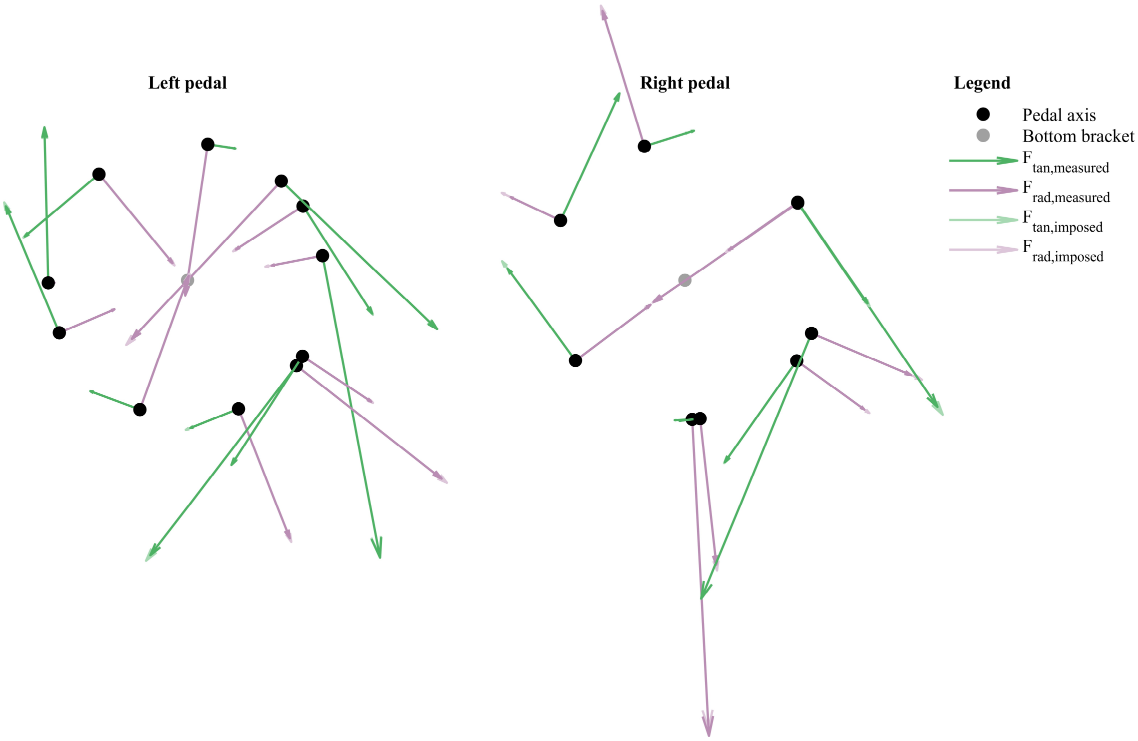


**Figure S1.** The measured and imposed tangential ($F_{tan}$) and radial ($F_{rad}$) pedal force vectors for each crank angle.

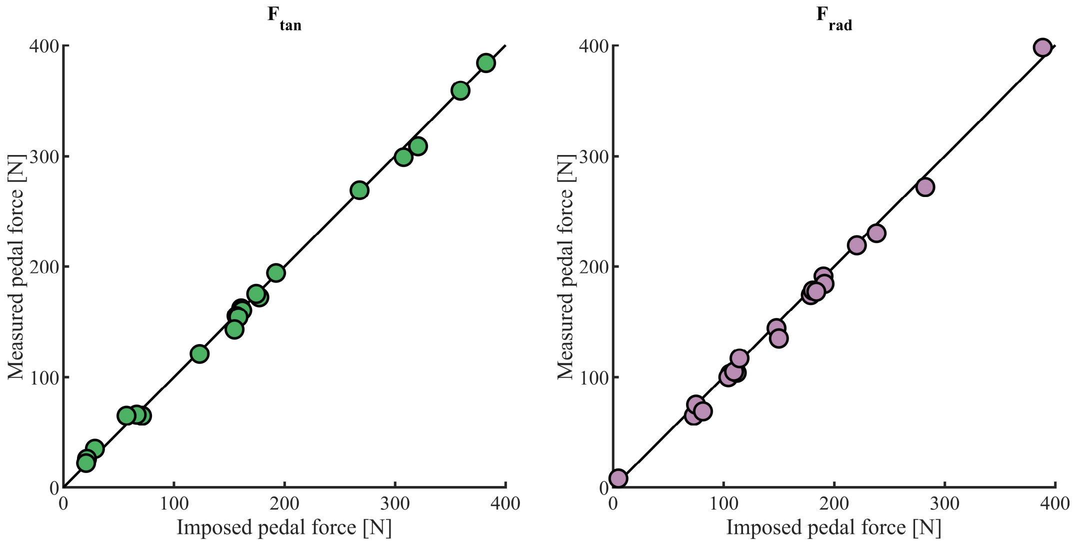


**Figure S2.** The measured tangential ($F_{tan}$) and radial ($F_{rad}$) pedal force magnitudes against the imposed pedal force magnitudes. The black line equals the line of identity.

*Dynamic validation – drag test on treadmill*

We performed a drag test to dynamically validate the power output measured by the pedals and crank. The protocol comprised two parts: 1) determining the imposed power output, and 2) determining the power output measured by the pedals and crank for each imposed power output. During both parts, one participant cycled on a treadmill (skate treadmill RL4500E; Rodby Innovation AB, Vänge, Sweden).

The imposed power output was determined by 'dragging' the participant, who sat on the bicycle without pedalling, using a rope that was attached to the front of the bicycle and to a force sensor (LSB201 S Beam Jr.® Load Sensor; Futek, Irvine, CA, USA), which was attached to a railing in front of the treadmill (see Fig. S3). We imposed four different power outputs by using two different belt speeds (8.4 and 10.0 m/s) and by adding two different weights (1.0 and 1.7 kg)

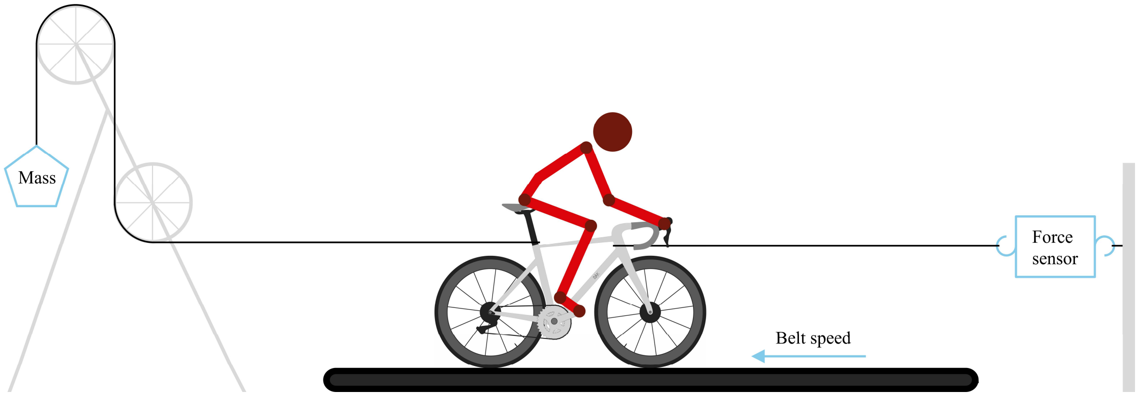


**Figure S3.** Schematic representation of the experimental setup for dynamically validating the power output measured by the pedals and crank.

to a rope that was attached to the rear of the bicycle and connected with a system of pulleys behind the treadmill (see Fig. S3). For each imposed power output, the force required to drag the participant was measured by the force sensor (at 100 Hz) for 1 min. The imposed power output equalled the measured average force multiplied with the belt speed.

We determined the power output measured by the pedals and crank for each imposed power output by removing the rope in front of the bicycle and having the participant cycle in each of the belt speed and added weight combinations. To investigate the impact of cadence, the participant cycled at five gear ratios: the preferred gear ratio, one and two gear ratios lower than preferred, and one and two gear ratios higher than preferred. The participant cycled for 30 seconds at each cadence. For each imposed power output and cadence, we calculated the average power output measured by the pedals and crank (at 200 Hz). We validated the measured with the imposed power output by calculating the bias, precision and RMSE.

Fig. S4 compares the measured and imposed power output, together with the different cadences used. The bias ± precision (RMSE) for the pedals and crank were -5 ± 7 W (8 W) and -16 ± 7 W (17 W), respectively. Cadence had no consistent effect on the measured power output. We concluded that the power output measured by the pedals shows good dynamic validity, whereas the power output measured by the crank considerably underestimates the imposed power output.

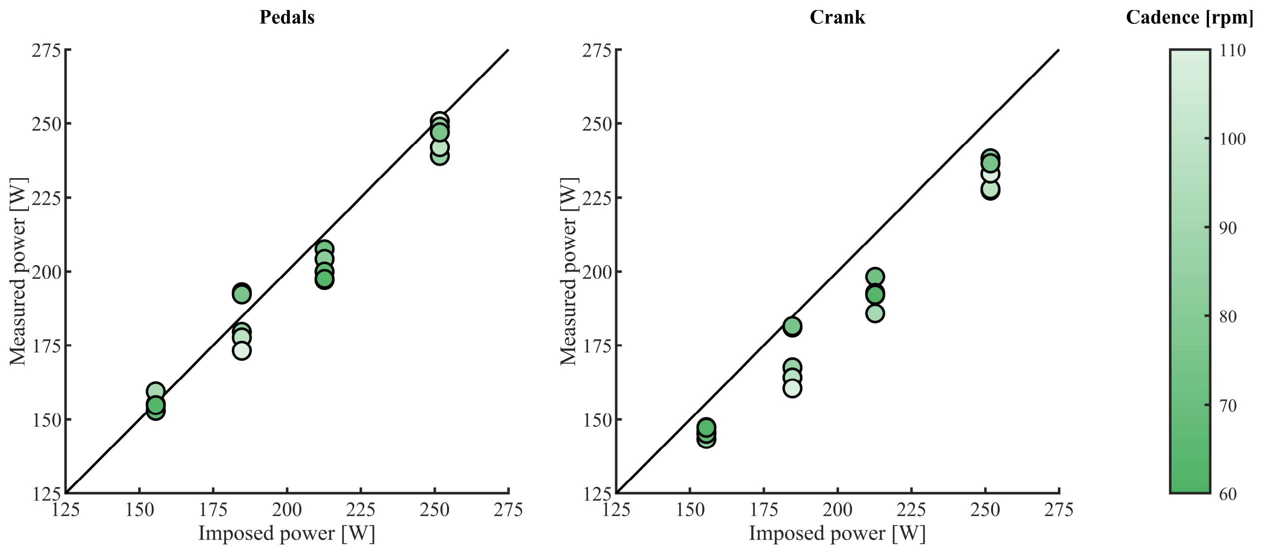


**Figure S4.** The measured power output by the pedals and crank against the imposed power output. The black line equals the line of identity.

*Dynamic validation – sprint test on ergometer*

The maximal average mechanical power output (AMPO) over one revolution reported in the main article was measured by the pedals. The data acquisition, however, was not limited to the pedals, as we simultaneously measured the mechanical power output using the crank and ergometer. In this validation, we determined the maximal AMPO as well as the maximal AMPO with 1 AFO and 2 AFOs relative to that without AFOs measured by the pedals, crank and ergometer. We compared these among the power meters by calculating the bias, precision and RMSE of the crank and ergometer relative to the pedals.

Fig. S5 compares the maximal AMPO of the crank and ergometer with that of the pedals. Relative to the pedals, the bias ± precision (RMSE) in maximal AMPO for the crank and ergometer were -64 ± 19 W (66 W) and -26 ± 24 W (35 W), respectively. Fig. S6 compares the relative maximal AMPO with 1 AFO and 2 AFOs of the crank and ergometer with that of the pedals. With 1 AFO, bias ± precision (RMSE) for the crank and ergometer were 1 ± 1% (1%) and -1 ± 3% (3%), respectively. With 2 AFOs, these were 1 ± 2% (2%) and -1 ± 2% (2%), respectively. Taken together, whereas the maximal AMPO of the crank and ergometer were lower than that of the pedals, the relative maximal AMPO with 1 AFO and 2 AFOs were similar to those of the pedals. Therefore, the impact of ankle immobility on sprint cycling performance was found to be similar across these power meters.

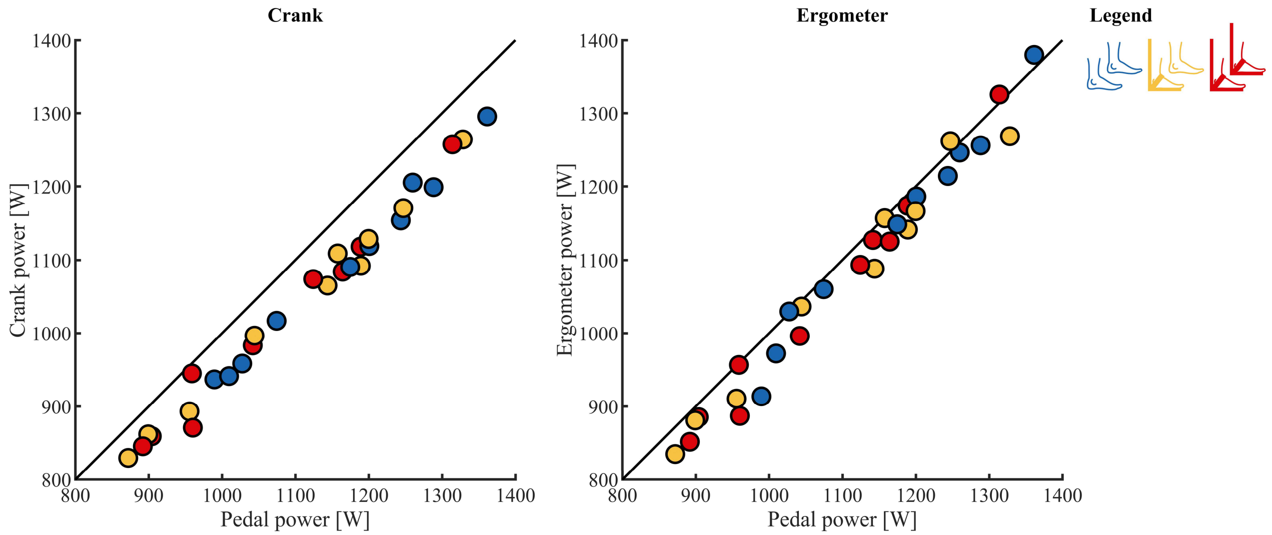


**Figure S5.** The measured power output by the crank and ergometer against that measured by the pedals. The black line equals the line of identity.

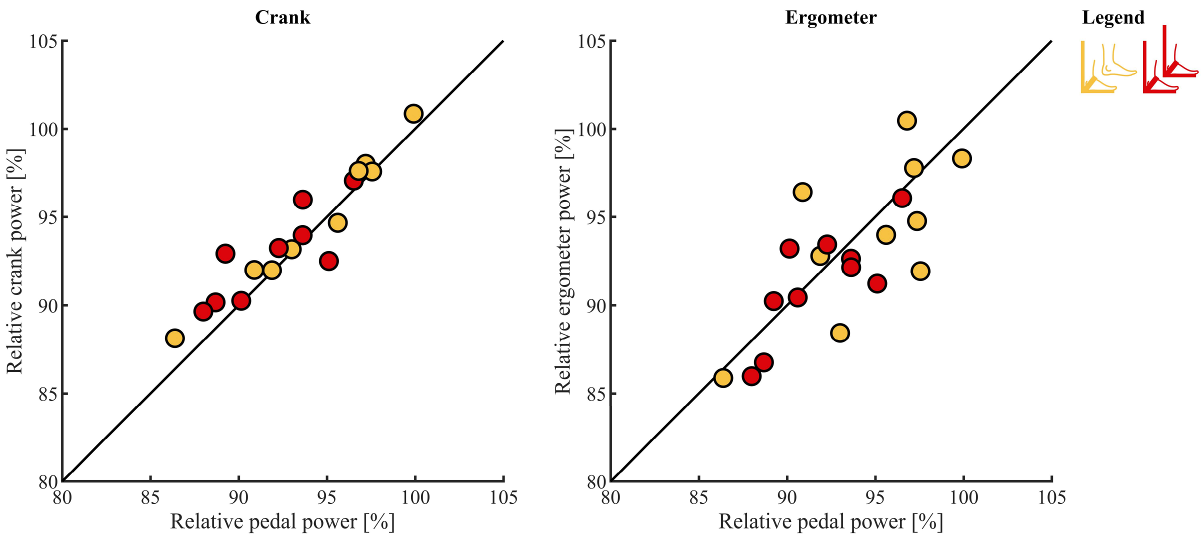


**Figure S6.** The measured power output by the crank and ergometer relative to that without AFOs against that measured by the pedals. The black line equals the line of identity. AFO, ankle foot orthosis.
